# Amyloid-β 42-carrying extracellular vesicles are associated with neurodegeneration and neuroinflammation in Alzheimer’s disease

**DOI:** 10.1101/2025.11.27.690912

**Authors:** Lukas Raich, Mirko Könken, Katarzyna M. Grochowska, Berta Puig, Christina Mayer, Alexandros Hadjilaou, Eckhard Schlemm, Mohsin Shafiq, Götz Thomalla, Tim Magnus, Jürgen Gallinat, Tharick A. Pascoal, Serge Gauthier, Markus Glatzel, Min Shi, Jing Zhang, Pedro Rosa-Neto, Marcel S. Woo

## Abstract

Alzheimer’s disease (AD) is the most common form of dementia, characterized by progressive amyloid-β (Aβ) accumulation and neurodegeneration. Extracellular vesicles (EVs) have been implicated in AD pathology, but their relationship to disease progression and the cerebrospinal fluid (CSF) proteome remains poorly understood. In this study, we analysed plasma-derived Aβ42-containing EVs from participants of the Alzheimer’s Disease Neuroimaging Initiative (ADNI) and calculated the ratio between the percentage of Aβ42-positive peripheral EVs and CSF Aβ42 (rAβ42). This ratio was elevated in AD and positively correlated with amyloid pathology as measured by PET. Longitudinal analyses revealed that higher rAβ42 values predicted accelerated hippocampal atrophy and cognitive decline, independent of CSF Aβ42 concentrations. Proteomic profiling of CSF using the SomaScan platform showed that rAβ42 was associated with an inflammatory signature characterized by myeloid activation and type II interferon-related signalling pathways. Together, these findings indicate that Aβ42-carrying EVs reflect an active and inflammatory component of AD pathology and may represent a novel class of biomarkers for disease progression. Understanding the mechanisms that link Aβ42-positive EVs to neuroinflammatory signalling could open new avenues for pathway-specific diagnostics and therapeutic strategies in AD.

## Introduction

Alzheimer’s disease (AD) is the most prevalent form of dementia worldwide and represents a major socio-economic burden ^1^. Its pathogenesis is characterized by the sequential accumulation of amyloid-β (Aβ) plaques and neurofibrillary tangles composed of hyperphosphorylated tau, ultimately leading to neurodegeneration and cognitive decline ^2^. In addition to these classical hallmarks, neuroinflammation has emerged as a key contributor to disease progression ^3^. Activated microglia and astrocytes can exert both detrimental and protective effects, on the one hand amplifying neurotoxicity and synaptic dysfunction ^4–6^, while on the other hand limiting Aβ deposition through phagocytic clearance ^7,8^. A deeper understanding of the distinct inflammatory mechanisms in AD may reveal novel biomarkers and therapeutic targets capable of predicting and modulating neurodegenerative processes.

The emergence of ultrasensitive protein detection technologies has revolutionized the biomarker landscape in neurodegenerative diseases. Highly sensitive cerebrospinal fluid (CSF) and blood assays now enable the detection ^9–12^ and staging ^13,14^ of pathophysiological processes across the full spectrum of Alzheimer’s disease, providing a more comprehensive view of disease progression ^15^. In particular, synaptic biomarkers have gained attention as strong predictors of cognitive decline in AD ^16,17^. However, despite these advances, there remains a critical need for additional biomarkers that not only detect disease-related pathologies but also predict clinical trajectories and individual rates of cognitive decline which might pave the way toward personalized diagnosis and therapy.

Extracellular vesicles (EVs) are lipid bilayer-enclosed particles that are actively secreted by virtually all cell types ^18^. They have gained increasing attention as biomarkers for central nervous system (CNS) disorders because they can cross the blood-brain barrier (BBB)^19^ and carry molecular cargo that reflects the identity and state of their cells of origin. Unlike most other biofluid biomarkers, EVs thus provide an opportunity to assess cell type-specific pathophysiological processes in vivo. Beyond their diagnostic potential, EVs are also biologically active mediators of intercellular communication. For instance, tau and its hyperphosphorylated forms have been detected within EVs ^20^, which is how they may facilitate the propagation of tau pathology ^21^. Furthermore, Aβ-oligomers can be detected in EVs isolated from the blood which correlate with AD hallmarks, and preclincial mechanistic studies showed that EVs from AD mouse models impair the neurovascular unit (NVU) ^22,23^. Moreover, EVs can modulate synaptic function ^24,25^ and immune cell activation in various disease contexts ^26^. Together, these properties make EVs promising biomarkers and mechanistic tools for understanding and monitoring AD progression.

In this study, we hypothesized that EVs containing Aβ42 oligomers are associated with disease progression in AD. We further examined whether these EVs are linked to distinct biological pathways by integrating CSF SomaScan proteomics data. Together, our findings uncover novel functions of Aβ42-containing EVs and provide insights into their potential role in mediating and reflecting AD pathophysiology.

## Results

### Cohort demographics

In this study, we included 100 cognitvely unimpaired (CU; 48 females) individuals, 92 people with mild cognitive impairment (MCI; 38 females) and 96 people with Alzheimer’s disease dementia (ADD; 42 females) from the Alzheimer’s Disease Neuroimaging Initiative (ADNI) where plasma EVs were isolated and the proportion of Aβ42-positive EVs was measured ^23^. The demographics are shown in **Table 1**. The participants could be separated according to the Aβ and tau brain load positivity (A/T) stages into 71 A-T-, 32 A+T-, 166 A+T+ and 19 A-T+ individuals. The EV data was only generated cross-sectionally for one timepoint. For all individuals longitudinal clinical, neuroimaging, and CSF biomarkers were available. The number of individuals with longitudinal follow-up per parameter is shown in the **Extended Data Table 1**.

**Table 1.** Cohort demographics.

| Diagnosis | Cognitively unimpaired | Mild cognitive impairment | Alzheimer's disease dementia |
| --- | --- | --- | --- |
| N (%) | 100 (34.7) | 92 (32.0) | 96 (33.33) |
| N female (%) | 48 (48) | 38 (41.3) | 42 (43.75) |
| APOE ε4 carriership, N(%) | 15 (15) | 66 (71.74) | 72 (75) |
| Age, years, mean (SD) | 74.04 (5.72) | 72.76 (7.32) | 75.16 (8.36) |
| Mini Mental State Exam, mean (SD) | 29.16 (1.12) | 26.87 (1.78) | 23.11 (1.97) |
| Clinical Dementia Rating Sum of Boxes, mean (SD) | 0.03 (0.12) | 1.95 (0.99) | 4.33 (1.57) |
| Neocortical [<sup>18</sup>F]Florbetapir, SUVR, mean (SD) | 1.03 (0.08) | 1.45 (0.17) | 1.41 (0.23) |
| CSF amyloid 1-42, pg/mL, mean (SD) | 1354.44 (369.33) | 659.64 (196.84) | 616.93 (277.83) |
| CSF p-tau181, pg/mL, mean (SD) | 17.91 (4.95) | 38.46 (14.05) | 39.28 (14.26) |
| CSF tau, pg/mL, mean (SD) | 202.28 (57.05) | 376.83 (125.2) | 389.95 (125.53) |
| Hippocampus volume, mm<sup>3</sup>, mean (SD) | 7466.86 (866.77) | 6281.27 (978.57) | 5725.72 (1093.47) |

### The Aβ42 EV / CSF Aβ42 ratio is increased in Alzheimer’s disease

First, we analysed the CSF Aβ42 and proportion of Aβ42-positive plasma EVs across the different diagnostic groups and A/T stages. As expected, since Aβ42 oligomers are sequestered in Aβ-plaques, we identified a significant decrease of Aβ42 in the diagnostic groups and A/T stages (**Fig. 1a**) while the proportion of Aβ42-positive plasma EVs (Aβ42+ EVs) was only significantly decreased in A+T+ in comparison to A-T- and did not change across the diagnostic groups (**Fig. 1b**). To account for the sequestered and inaccessible Aβ42 in the CNS in Aβ-plaques, we next normalized the proportion of Aβ42-positive plasma EVs to the CSF Aβ42 (rAβ42). This allowed us to investigate the effect of Aβ42-carrying EVs relative to the soluble Aβ42 that can be released from the CNS. This ratio increased in MCI and ADD in comparison to CU as well as in A+T+ in comparison to A-T- (**Fig. 1c**). To assess the correlation with cerebral Aβ, we performed region-wise correlation analysis with [^18^F]Florbetapir-measured Aβ-load. As expected, we found strong negative associations between CSF Aβ42 and Aβ-load in cortical regions in A+ and all individuals (**Fig. 1d**). While the Aβ42+ EVs were positively associated with Aβ-load in the pre- and postcentral gyri in A+ (**Fig. 1e**), rAβ42 was positively associated with Aβ-load in numerous cortical regions in A+ and across all participants (**Fig. 1d**). Importantly, this was still the case after including CSF Aβ42 as covariate (**Extended Data Fig. 1**), underlining that the positive association between rAβ42 and cortical Aβ-load is independent of the CSF Aβ42 levels.

**Fig. 1.**
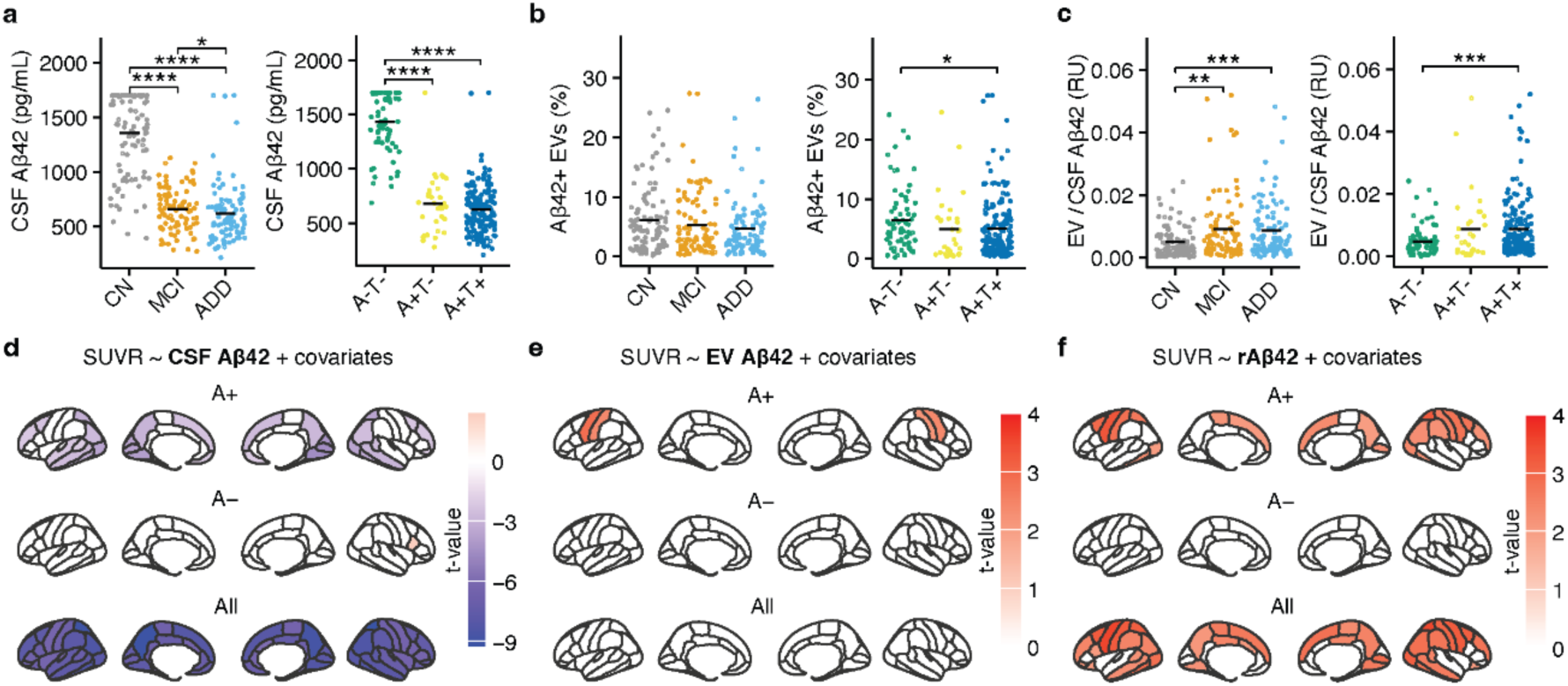
The ratio between Aβ42-positive extracellular vesicles and CSF Aβ42 is increased in Alzheimer’s disease. **(a-c)** CSF Aβ42 (pg/mL; a), the proportion of Aβ42-positive plasma extracellular vesicles (EVs; b), and the ratio between Aβ42-positive EVS and CSF Aβ42 (rAβ42; c) in cognitively unimpaired people (CU, n = 100), people with mild cognitive impairment (MCI, n = 92), and Alzheimer’s disease dementia (ADD, n = 96) or across the A/T stages (A-T-, n = 71; A+T-, n = 32; A+T+, n = 166). Mann-Whitney-U tests with FDR-correction were used. **(d-f)** Region-wise linear models between [^18^F]Florbetapir and CSF Aβ42 (d), the proportion of Aβ42-positive EVs (e), and rAβ42 (f). The Desikan-Killiany-Tourville cortical parcellation was used. Age, sex at birth and APOE4 carriership were used as covariates. T-values are only shown where the FDR-corrected *P*-values reached statistical significance *P* < 0.05. \**P* < 0.05, \*\**P* < 0.01, \*\*\**P* < 0.001, \*\*\*\**P* < 0.0001.

To further investigate the impact of Aβ42-carrying plasma EVs, we separated our cohort into rAβ42-high as defined by the top 25%, and low participants as defined by the lower 75%. Since the proportion of Aβ42-carrying plasma EVs and CSF Aβ42 were not significantly correlated (**Fig. 2a**), the rAβ42-groups represent a distinct biological aspect of AD. First, we compared the baseline characteristics of rAβ42-high vs. low in A- and A+ (A- rAβ42 low, n = 56; A+ rAβ42 low, n = 104; A- rAβ42 high, n = 7; A+ rAβ42 high, n = 45). As expected, the Aβ42+ EVs were increased (**Fig. 2b**) and the CSF Aβ42 (**Fig. 2c**) was decreased in rAβ42-high individuals independent of the amyloid-status underlining that the rAβ42 is not solely driven by either CSF Aβ42 decrease or an increase of the Aβ42-carrying plasma EVs. In contrast, the neocortical Aβ-load measured by [^18^F]Florbetapir (**Fig. 2d**), plasma p-tau181 (**Fig. 2e**), the hippocampus volume (**Fig. 2f**) as well as the cognitive scores (**Extended Data Fig. 2a-b**) were similar between these groups. The CSF p-tau181 and total tau were significantly lower in the A- rAβ42 high individuals while no differences between the rAβ42 groups were observed in A+ individuals (**Extended Data Fig. 2c-d**).

**Fig. 2.**
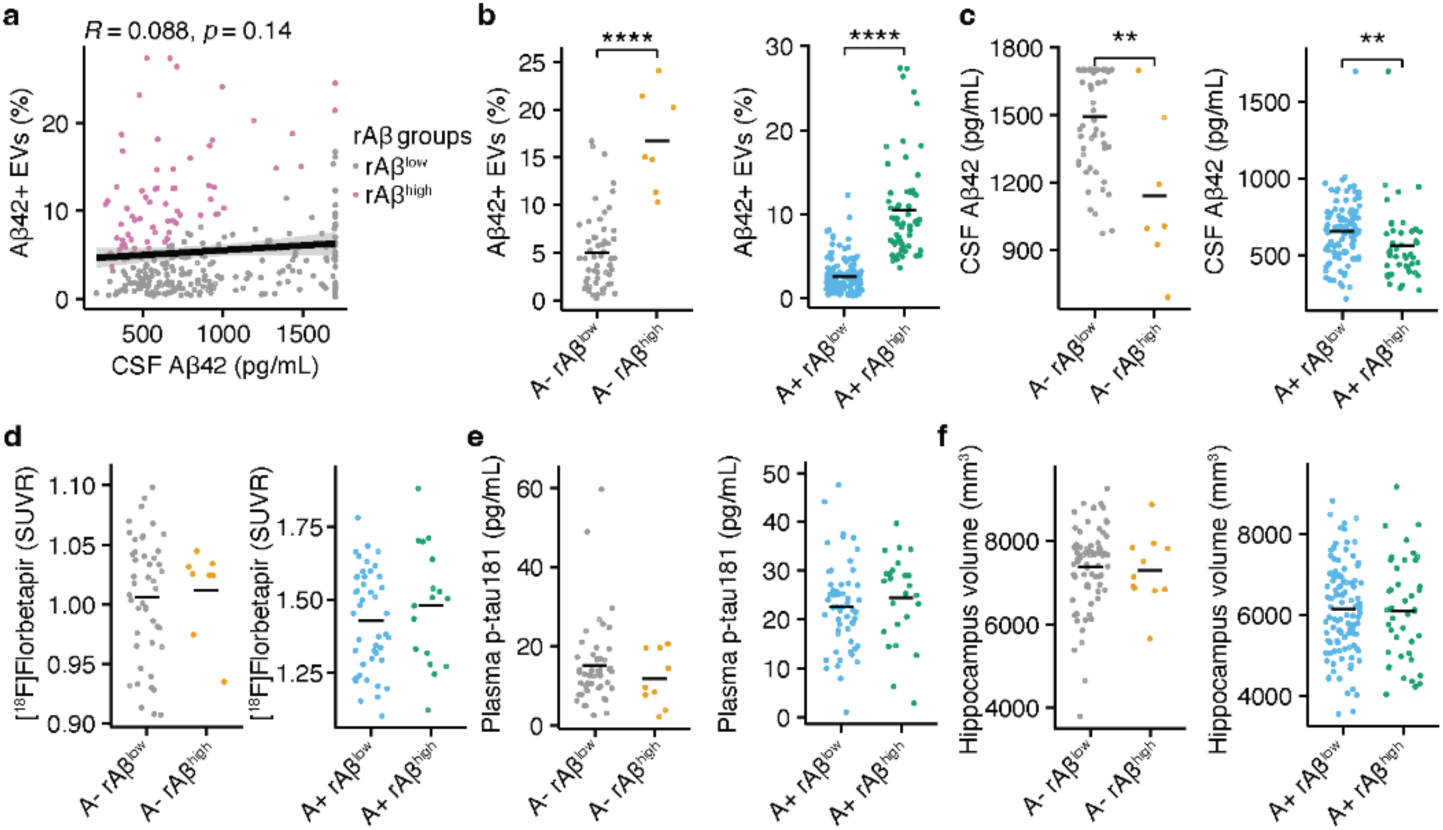
The EV/CSF Aβ42-ratio captures a distinct AD disease state. **(a)** Spearman correlation analysis between the % of Aβ42-positive plasma extracellular vesicles (EVs) and CSF Aβ42. The ratio between Aβ42-positive plasma EVS and CSF Aβ42 (rAβ42) was used to identify the 25% participants with the highest rAβ42 (rAβ42 high) which are labeled in pink in the plot. **(b-f)** % of Aβ42-positive plasma EVs (b), CSF Aβ42 (c), neocortical [^18^F]Florbetapir standard uptake value ratio (SUVR; d), plasma p-tau181 (e) and hippocampus volume (f) in rAβ42 high vs. rAβ42 low people with (A+) or without (A-) Aβ pathology (A- rAβ42 low, n = 56; A+ rAβ42 low, n = 104; A- rAβ42 high, n = 7; A+ rAβ42 high, n = 45). rAβ42 = ratio between the % of Aβ42-positive plasma EVs and CSF Aβ42. Mann-Whitney-U tests were used. \*\**P* < 0.01, \*\*\*\**P* < 0.0001.

### Aβ42-containing EVs predict neurodegeneration and cognitive decline

In the next step, we aimed to investigate how Aβ42-carrying plasma EVs influence the longitudinal AD trajectory. Therefore, we compared the longitudinal trajectories of rAβ42 high and low individuals with or without positive amyloid-status. The exact number of individuals and follow-up per group and biomarker are provided in the **Extended Data Table 1**. In A-, the longitudinal trajectories of neocortical [^18^F]Florbetapir, CSF p-tau181, CSF total tau, hippocampus volume and cognitive decline were statistically not different between rAβ42-high and low individuals (**Fig. 3a-c**, **Extended Data Fig. 3a-c**). In A+, neocortical [^18^F]Florbetapir, CSF p-tau181, CSF total tau significantly increased over time as expected but no differences were detected between rAβ42-high and low individuals (**Extended Data Fig. 3d-e**). In contrast, A+ rAβ42-high individuals showed faster neurodegeneration measured by hippocampus volume loss (**Fig. 3d**; Standardized β = -0.09, *P* = 0.01) and cognitive decline assessed by MMSE (**Fig. 3e**; Standardized β = -0.21, *P* = p0.01) and CDR-SOB (**Fig. 3f**; Standardized β = 0.17, *P* = 0.03). The full model statistics testing the interactions of the rAβ42-groups and longitudinal time as well as all covariates are shown in **Table 2** for the A- and in **Table 3** for the A+ individuals. The statistical results testing the longitudinal trajectories of A-/A+ rAβ42-high/low individuals separately is provided in the **Extended Data Table 2**.

**Fig. 3.**
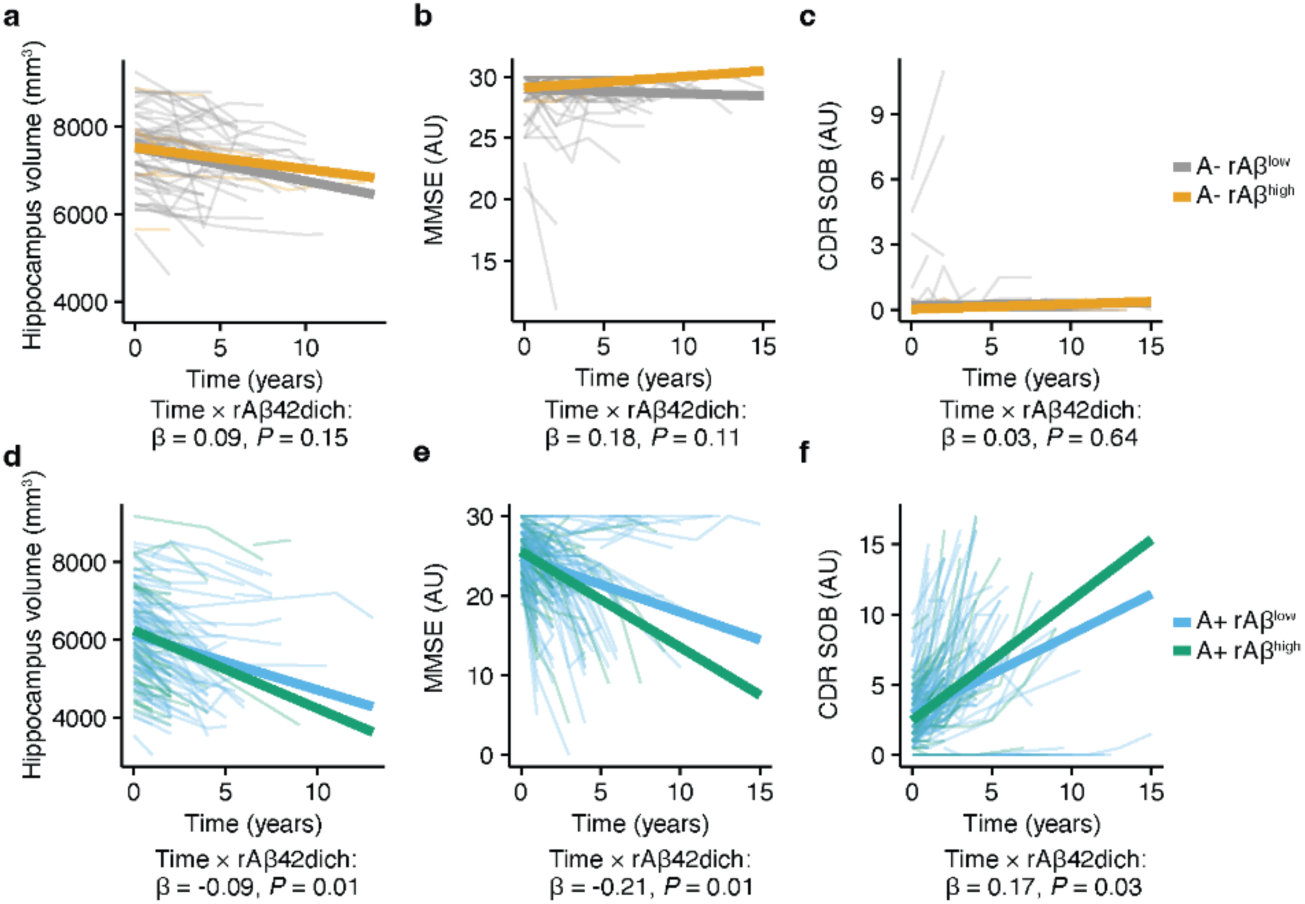
EV/CSF Aβ42-ratio predicts neurodegeneration and cognitive decline. **(a-c)** Longitudinal analysis of hippocampus volume (a), mini-mental state examination (MMSE) (b) and clinical dementia rating sum of boxes (CDR-SOB) (c) in rAβ42 high and rAβ42 low participants without Aβ pathology. The standardized β and *P*-value for the interaction between time and rAβ42-status are shown in the figure. **(d-f)** Longitudinal analysis of hippocampus volume (d), MMSE (e) and CDR-SOB (f) in rAβ42 high and rAβ42 low participants with Aβ pathology. The standardized β and *P*-value for the interaction between time and rAβ42-status are shown in the figure. Age, sex at birth, APOE4 carriership, CSF Aβ42 baseline and CSF p-tau181 baseline values were used as covariates. The respective numbers of individuals for the different groups are shown in the **Extended Data Table 1**. The full statistical results for the interaction analysis between the time and rAβ42-status are shown in **Table 2 and 3**. Statistical results of the longitudinal biomarker progression in the rAβ42 high and rAβ42 low are shown in the **Extended Data Table 2**. rAβ42 = ratio between the % of Aβ42-positive plasma EVs and CSF Aβ42.

**Table 2.** Statistical results of longitudinal analyses in A- individuals.

| Amyloid status | Parameter | Term | Standardized $\beta$ (95% CIs) | P-value |
| --- | --- | --- | --- | --- |
| A- | CDR-SOB | Time | 0.03 (-0.02, 0.07) | 0.245 |
| A- | CDR-SOB | rAβ42 high | -0.16 (-0.33, 0.02) | 0.075 |
| A- | CDR-SOB | Age | 0.01 (-0.04, 0.06) | 0.695 |
| A- | CDR-SOB | Sex at birth | 0.06 (-0.04, 0.17) | 0.224 |
| A- | CDR-SOB | APOE4 carriership | -0.01 (-0.06, 0.04) | 0.754 |
| A- | CDR-SOB | Baseline | 0.97 (0.9, 1.03) | <0.001 |
| A- | CDR-SOB | CSF Aβ42 baseline | < 0.001 (-0.06, 0.05) | 0.904 |
| A- | CDR-SOB | CSF p-tau181 baseline | -0.06 (-0.14, 0.01) | 0.075 |
| A- | CDR-SOB | Time x rAβ42 groups | 0.03 (-0.1, 0.16) | 0.641 |
| A- | Hippocampus volume | Time | -0.25 (-0.3, -0.21) | <0.001 |
| A- | Hippocampus volume | rAβ42 high | 0.03 (-0.14, 0.21) | 0.652 |
| A- | Hippocampus volume | Age | 0.02 (-0.04, 0.08) | 0.539 |
| A- | Hippocampus volume | Sex at birth | -0.03 (-0.13, 0.08) | 0.624 |
| A- | Hippocampus volume | APOE4 carriership | 0.04 (-0.01, 0.1) | 0.09 |
| A- | Hippocampus volume | Baseline | 0.95 (0.89, 1.01) | <0.001 |
| A- | Hippocampus volume | CSF Aβ42 baseline | -0.04 (-0.09, 0.02) | 0.189 |
| A- | Hippocampus volume | CSF p-tau181 baseline | -0.02 (-0.07, 0.04) | 0.543 |
| A- | Hippocampus volume | Time x rAβ42 groups | 0.09 (-0.03, 0.2) | 0.149 |
| A- | MMSE | Time | -0.06 (-0.14, 0.01) | 0.094 |
| A- | MMSE | rAβ42 high | 0.31 (-0.02, 0.64) | 0.484 |
| A- | MMSE | Age | -0.08 (-0.18, 0.02) | 0.123 |
| A- | MMSE | Sex at birth | -0.11 (-0.31, 0.08) | 0.262 |
| A- | MMSE | APOE4 carriership | 0.1 (0, 0.2) | 0.063 |
| A- | MMSE | Baseline | 0.7 (0.59, 0.81) | <0.001 |
| A- | MMSE | CSF Aβ42 baseline | 0.13 (0.03, 0.23) | 0.015 |
| A- | MMSE | CSF p-tau181 baseline | -0.14 (-0.25, -0.03) | 0.013 |
| A- | MMSE | Time x rAβ42 groups | 0.18 (-0.04, 0.4) | 0.108 |
| A- | CSF p-tau 181 | Time | 0.08 (0.06, 0.11) | <0.001 |
| A- | CSF p-tau 181 | rAβ42 high | 0.1 (-0.02, 0.22) | 0.393 |
| A- | CSF p-tau 181 | Age | 0.01 (-0.02, 0.05) | 0.411 |
| A- | CSF p-tau 181 | Sex at birth | -0.04 (-0.1, 0.03) | 0.285 |
| A- | CSF p-tau 181 | APOE4 carriership | 0.02 (-0.01, 0.05) | 0.257 |
| A- | CSF p-tau 181 | Baseline | 0.99 (0.96, 1.01) | <0.001 |
| A- | CSF p-tau 181 | CSF Aβ42 baseline | 0.03 (0, 0.06) | 0.087 |
| A- | CSF p-tau 181 | Time x rAβ42 groups | 0.05 (-0.06, 0.16) | 0.355 |
| A- | CSF tau | Time | 0.1 (0.07, 0.13) | <0.001 |
| A- | CSF tau | rAβ42 high | 0.03 (-0.13, 0.19) | 0.965 |
| A- | CSF tau | Age | 0.02 (-0.03, 0.07) | 0.484 |
| A- | CSF tau | Sex at birth | -0.09 (-0.17, 0) | 0.053 |
| A- | CSF tau | APOE4 carriership | 0.02 (-0.02, 0.07) | 0.331 |
| A- | CSF tau | Baseline | 1.01 (0.76, 1.26) | <0.001 |
| A- | CSF tau | CSF Aβ42 baseline | 0.01 (-0.04, 0.06) | 0.759 |
| A- | CSF tau | CSF p-tau181 baseline | -0.03 (-0.27, 0.2) | 0.774 |
| A- | CSF tau | Time x rAβ42 groups | 0.03 (-0.11, 0.18) | 0.677 |
| A- | [18F]Florbetapir | Time | 0.04 (-0.04, 0.12) | 0.35 |
| A- | [18F]Florbetapir | rAβ42 high | -0.14 (-0.5, 0.21) | 0.54 |
| A- | [18F]Florbetapir | Age | 0.01 (-0.11, 0.14) | 0.817 |
| A- | [18F]Florbetapir | Sex at birth | 0.03 (-0.22, 0.27) | 0.838 |
| A- | [18F]Florbetapir | APOE4 carriership | 0.06 (-0.05, 0.17) | 0.311 |
| A- | [18F]Florbetapir | Baseline | 0.82 (0.7, 0.94) | <0.001 |
| A- | [18F]Florbetapir | CSF Aβ42 baseline | -0.05 (-0.18, 0.07) | 0.378 |
| A- | [18F]Florbetapir | CSF p-tau181 baseline | -0.01 (-0.12, 0.09) | 0.795 |
| A- | [18F]Florbetapir | Time x rAβ42 groups | -0.26 (-0.52, 0) | 0.051 |

**Table 3.** Statistical results of longitudinal analyses in A+ individuals.

| Amyloid status | Parameter | Term | Standardized $\beta$ (95% CIs) | P-value |
| --- | --- | --- | --- | --- |
| A+ | CDR-SOB | Time | 0.47 (0.4, 0.54) | <0.001 |
| A+ | CDR-SOB | rAβ42 high | -0.11 (-0.26, 0.04) | 0.006 |
| A+ | CDR-SOB | Age | 0.01 (-0.06, 0.08) | 0.745 |
| A+ | CDR-SOB | Sex at birth | -0.06 (-0.2, 0.07) | 0.352 |
| A+ | CDR-SOB | APOE4 carriership | 0.01 (-0.06, 0.08) | 0.806 |
| A+ | CDR-SOB | Baseline | 0.64 (0.57, 0.72) | <0.001 |
| A+ | CDR-SOB | CSF Aβ42 baseline | -0.26 (-0.34, -0.18) | <0.001 |
| A+ | CDR-SOB | CSF p-tau181 baseline | 0.1 (0.04, 0.17) | 0.003 |
| A+ | CDR-SOB | Time x rAβ42 groups | 0.17 (0.02, 0.33) | 0.029 |
| A+ | Hippocampus volume | Time | -0.27 (-0.3, -0.24) | <0.001 |
| A+ | Hippocampus volume | rAβ42 high | 0.03 (-0.05, 0.1) | 0.016 |
| A+ | Hippocampus volume | Age | 0.01 (-0.02, 0.05) | 0.453 |
| A+ | Hippocampus volume | Sex at birth | 0.07 (0, 0.14) | 0.044 |
| A+ | Hippocampus volume | APOE4 carriership | -0.01 (-0.05, 0.02) | 0.541 |
| A+ | Hippocampus volume | Baseline | 0.96 (0.92, 1) | <0.001 |
| A+ | Hippocampus volume | CSF Aβ42 baseline | 0.06 (0.02, 0.09) | 0.004 |
| A+ | Hippocampus volume | CSF p-tau181 baseline | -0.04 (-0.07, -0.01) | 0.011 |
| A+ | Hippocampus volume | Time x rAβ42 groups | -0.09 (-0.17, -0.02) | 0.012 |
| A+ | MMSE | Time | -0.41 (-0.48, -0.33) | <0.001 |
| A+ | MMSE | rAβ42 high | 0.07 (-0.09, 0.24) | 0.012 |
| A+ | MMSE | Age | < 0.001 (-0.08, 0.07) | 0.923 |
| A+ | MMSE | Sex at birth | 0.11 (-0.04, 0.25) | 0.156 |
| A+ | MMSE | APOE4 carriership | -0.03 (-0.11, 0.05) | 0.433 |
| A+ | MMSE | Baseline | 0.58 (0.51, 0.66) | <0.001 |
| A+ | MMSE | CSF Aβ42 baseline | 0.26 (0.18, 0.35) | <0.001 |
| A+ | MMSE | CSF p-tau181 baseline | -0.14 (-0.22, -0.07) | <0.001 |
| A+ | MMSE | Time x rAβ42 groups | -0.21 (-0.37, -0.05) | 0.012 |
| A+ | CSF p-tau 181 | Time | 0.05 (0.03, 0.08) | <0.001 |
| A+ | CSF p-tau 181 | rAβ42 high | < 0.001 (-0.07, 0.07) | 0.338 |
| A+ | CSF p-tau 181 | Age | < 0.001 (-0.03, 0.03) | 0.935 |
| A+ | CSF p-tau 181 | Sex at birth | 0.03 (-0.03, 0.09) | 0.355 |
| A+ | CSF p-tau 181 | APOE4 carriership | -0.01 (-0.04, 0.03) | 0.722 |
| A+ | CSF p-tau 181 | Baseline | 0.96 (0.93, 0.99) | <0.001 |
| A+ | CSF p-tau 181 | CSF Aβ42 baseline | 0.01 (-0.02, 0.05) | 0.447 |
| A+ | CSF p-tau 181 | Time x rAβ42 groups | -0.05 (-0.11, 0.01) | 0.112 |
| A+ | CSF tau | Time | 0.11 (0.08, 0.14) | <0.001 |
| A+ | CSF tau | rAβ42 high | -0.01 (-0.08, 0.06) | 0.785 |
| A+ | CSF tau | Age | 0.01 (-0.02, 0.05) | 0.424 |
| A+ | CSF tau | Sex at birth | -0.01 (-0.07, 0.06) | 0.785 |
| A+ | CSF tau | APOE4 carriership | -0.02 (-0.05, 0.02) | 0.375 |
| A+ | CSF tau | Baseline | 0.93 (0.75, 1.11) | <0.001 |
| A+ | CSF tau | CSF Aβ42 baseline | -0.02 (-0.05, 0.02) | 0.37 |
| A+ | CSF tau | CSF p-tau181 baseline | 0.03 (-0.15, 0.21) | 0.772 |
| A+ | CSF tau | Time x rAβ42 groups | -0.03 (-0.09, 0.04) | 0.415 |
| A+ | [18F]Florbetapir | Time | 0.15 (0.08, 0.23) | < 0.001 |
| A+ | [18F]Florbetapir | rAβ42 high | -0.09 (-0.26, 0.08) | 0.695 |
| A+ | [18F]Florbetapir | Age | < 0.001 (-0.07, 0.07) | 1 |
| A+ | [18F]Florbetapir | Sex at birth | -0.06 (-0.2, 0.09) | 0.453 |
| A+ | [18F]Florbetapir | APOE4 carriership | -0.03 (-0.11, 0.05) | 0.409 |
| A+ | [18F]Florbetapir | Baseline | 0.91 (0.82, 0.99) | < 0.001 |
| A+ | [18F]Florbetapir | CSF Aβ42 baseline | -0.08 (-0.16, 0.01) | 0.086 |
| A+ | [18F]Florbetapir | CSF p-tau181 baseline | 0.02 (-0.05, 0.1) | 0.543 |
| A+ | [18F]Florbetapir | Time x rAβ42 groups | -0.05 (-0.22, 0.12) | 0.54 |

### The Aβ42 EV / CSF Aβ42 ratio is associated with neuroinflammation

Subsequently, we investigated which biological processes were associated with rAβ42. Therefore, we selected 145 individuals with amyloid pathology and 61 individuals without amyloid pathology where CSF SomaScan data was available. We performed linear regression models of rAβ42 with each protein adjusting for age, and sex at birth as covariates (**Fig. 4a** shows the results for A+, **Extended Data Fig. 4** shows the results for A-, all statistical results are shown in **Extended Data Table 3**). To identify biological pathways, we selected the CSF proteins with a positive association reaching statistical significance and performed a gene ontology (GO) biological process analysis. While we found no enriched GO terms for the A-individuals, inflammatory processes like the type II interferon response, myeloid activation or the tumor necrosis factor response were significantly enriched in the A+ individuals (**Fig. 4b**). Finally, as previously shown ^27^, we selected the type II interferon response and generated a multi-protein composite score. This score showed a significantly positive correlation with the rAβ42 in the A+ participants (**Fig. 4c**) underlining that Aβ42-carrying EVs are associated with neuroinflammatory processes in AD.

**Fig. 4.**
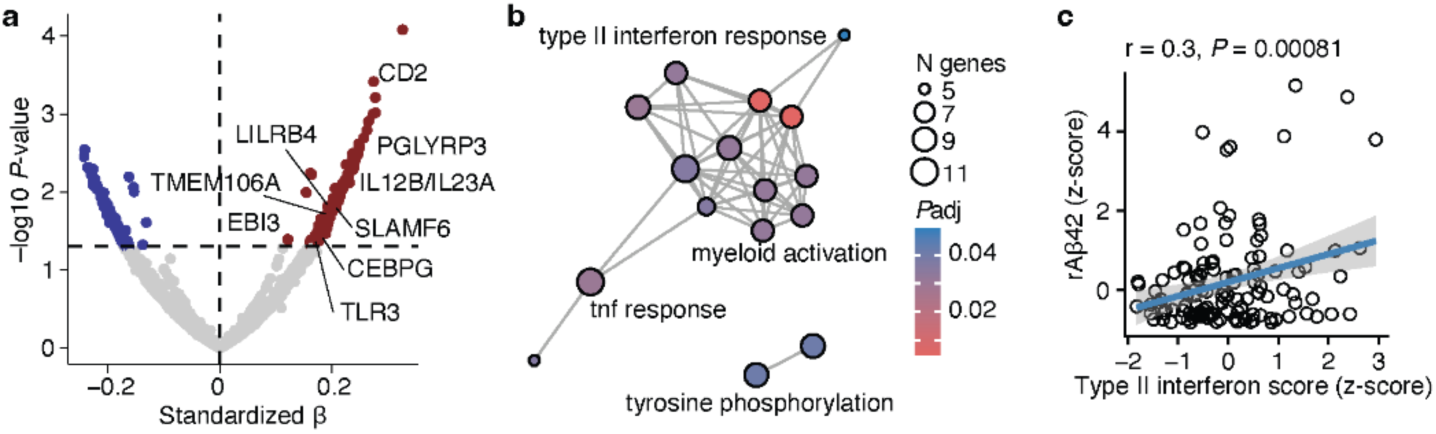
Aβ42-carrying extracellular vesicles are associated with neuroinflammation. **(a)** CSF 7K SomaScan data of 145 participants with amyloid pathology and available plasma EV and CSF Aβ42 was analyzed. A linear model testing the association between rAβ42 and each protein was used with age and sex at birth as covariates. The standardized β and the *P*-values are shown. Red and blue colors indicate significant up- or downregulation. **(b)** Results of gene ontology biological process analysis of the CSF proteins that were significantly positively associated with rAβ42. **(c)** Spearman correlation analysis between rAβ42 and a type II interferon composite score calculated from the 7K SomaScan data in participants with Aβ pathology (n = 145). rAβ42 = ratio between the % of Aβ42-positive plasma EVs and CSF Aβ42.

## Discussion

In this study, we demonstrated that EVs carrying Aβ42 predict hippocampal volume loss and cognitive decline in individuals with established amyloid pathology. Moreover, these Aβ42-positive EVs were associated with a distinct CSF proteomic signature enriched for inflammation-related pathways. Together, these findings provide novel insights into the interplay between amyloid pathology, neuroinflammation, and EV biology in Alzheimer’s disease, suggesting that Aβ42-containing EVs may serve as both biomarkers and mechanistic mediators of disease progression.

To investigate Aβ42-containing EVs, we analysed EVs isolated from plasma and calculated the ratio between the proportion of Aβ42-positive EVs and Aβ42 concentrations measured in the CSF. Although recent studies have reported that Aβ42-containing EVs decrease in AD ^23^, we observed an increase in this rAβ42 ratio. This finding highlights the potential utility of biomarker ratios, which can provide normalized measures of compartmental distribution and have proven valuable for other amyloid-associated ^12,28,29^ and, more recently, synaptic biomarkers ^16^. Notably, the rAβ42 was significantly associated with PET-measured amyloid accumulation across multiple cortical regions, even after adjusting for CSF Aβ42 levels. This suggests that Aβ42-carrying EVs exert an effect on cerebral amyloid deposition that is independent of total soluble Aβ42 concentrations in the CSF. Such findings support the notion that EV-bound Aβ42 may represent a distinct biological pathway contributing to amyloid aggregation and propagation within the brain which needs further mechanistic investigations. Next, we stratified participants into rAβ42-high and rAβ42-low groups and compared classical AD biomarkers across these subgroups. As expected, individuals with amyloid pathology and high rAβ42 exhibited lower CSF Aβ42 and a higher proportion of Aβ42-positive EVs, whereas total tau, p-tau181, cognitive status, and hippocampal volume did not differ significantly. Remarkably, neocortical Aβ load measured by [^18^F]Florbetapir-PET were similar between these groups, indicating that Aβ42-containing EVs might capture aspects of disease progression that are not reflected by PET-measured amyloid burden. Interestingly, among A-individuals, higher rAβ42 levels were associated with lower tau and p-tau181 CSF levels in our cross-sectional analyses and with a numerically slower increase of amyloid accumulation as measured by [^18^F]Florbetapir PET in comparison to A- rAβ-low individuals. Although, this did not reach statistical significance, this observation raises the possibility that in A-individuals, EVs could exert a protective role by facilitating Aβ clearance ^30^. However, the sample size of the A- participants in our cohort was limited, and these results should therefore be interpreted with caution. Future studies in larger, well-characterized cohorts are warranted to clarify the role of Aβ42-carrying EVs in early, preclinical stages of disease.

Conversely, rAβ42-high individuals with proven amyloid pathology exhibited faster neurodegeneration and cognitive decline than rAβ42-low individuals, as reflected by longitudinal hippocampal volume loss, decreases in MMSE scores and increases in CDR-SOB. However, we did not observe significant differences in the longitudinal trajectories of total tau, p-tau181, or PET-measured neocortical amyloid accumulation between the rAβ42 subgroups. Together, these findings underscore that EVs are associated with disease progression in AD. Previous studies have demonstrated that EVs containing APP fragments, Aβ peptides, or tau and its phosphorylated forms can induce synaptic dysfunction, although the underlying mechanisms remain incompletely understood ^22,31,32^. Additionally, alterations in the lipid composition of brain-derived EVs have been reported in AD ^33,34^, and EV-associated insulin-signalling markers have been shown to predict cognitive decline ^35^. Moreover, tau filaments have been detected tethered within EVs isolated from AD brains, supporting their potential role in the intercellular propagation of tau pathology ^20,21^. Although tau-PET data were not available for the participants included in this analysis, future studies combining tau-PET imaging with rAβ42 measures may provide deeper insight into the relationship between EV-mediated amyloid processes and tau pathology.

To identify biological pathways that might be associated with Aβ42-carrying EVs, we analysed CSF proteomics data obtained using the SomaScan platform. Such ultrasensitive proteomic technologies provide powerful tools to investigate disease-related signalling networks in vivo ^27,36–39^. In individuals with confirmed amyloid pathology, proteins linked to neuroinflammatory signalling, particularly myeloid activation and type II interferon-related inflammatory responses were significantly associated with Aβ42-containing EVs. Myeloid cell activation is a well-established contributor to AD pathogenesis ^5,40–42^, as well as interferon-driven responses which have also been implicated in neuronal injury and neurodegeneration in AD ^43^ and across neuroinflammatory and neurodegenerative disorders ^44,45^. Intriguingly, bidirectional communication between EVs and myeloid cells may amplify these processes. EVs can modulate myeloid activation states, and myeloid cells themselves release EVs that might influence neuronal synaptic function ^46–48^, as demonstrated in other disease contexts like ischemic stroke ^49,50^. Further preclinical studies are required to determine how Aβ42-containing EVs trigger neuroinflammatory stress responses across different brain-resident cell types. Such mechanistic insights may ultimately pave the way toward individualized, pathway-specific biomarkers and therapeutic strategies ^51^.

This study has several limitations. First, it was conducted as a monocentric study and thus requires validation in independent cohorts, particularly using distinct EV isolation and characterization methods. Second, we analysed whole EVs isolated from plasma, which may not fully capture CNS-intrinsic processes. Emerging single-EV and cell type-specific profiling technologies will enable a more precise understanding of cell-of-origin mechanisms in future work. Third, tau-PET data were not available in our cohort, preventing an assessment of the relationship between Aβ42-carrying EVs and tau propagation. Fourth, the number of A-participants was limited, restricting our ability to evaluate the role of Aβ42-positive EVs in individuals without established amyloid pathology which should be addressed in subsequent studies. Finally, while CSF proteomics revealed inflammatory pathways associated with Aβ42-carrying EVs, our findings remain correlative, and mechanistic studies are needed to determine causality.

In summary, our study demonstrates that Aβ42-carrying EVs are associated with longitudinal neurodegeneration and cognitive decline, as well as with an inflammatory CSF proteomic profile. These results highlight the potential of EVs as both biomarkers and mediators of AD progression, possibly leading to future biomarker-guided and pathway-specific therapeutic approaches.

## Methods

### Study population

We included cognitively unimpaired individuals, and MCI and ADD subjects from the Alzheimer’s Disease Neuroimaging Initiative (ADNI) database (adni.loni.usc.edu). The ADNI was launched in 2003 as a public-private partnership, led by Principal Investigator Michael W. Weiner, MD. The primary goal of ADNI has been to test whether serial magnetic resonance imaging (MRI), PET, other biological markers, and clinical and neuropsychological assessment can be combined to measure the progression of MCI and ADD. For up-to-date information, see www.adni-info.org. The study was approved by the Institutional Review Boards of all the participating institutions and informed written consent was obtained from all participants. Data used for the analyses presented here was accessed on August 1st, 2025. We included participants for whom Aβ1-42 in EVs, and [^18^F]Florbetapir PET or CSF Aβ1-42, p-tau181 in the CSF, and 7K SomaScan data was available. In ADNI, lumbar puncture was performed as described in the ADNI procedures manual (http://www.adni-info.org/). CSF samples were frozen on dry ice within 1 hour after collection and shipped overnight on dry ice to the ADNI Biomarker Core laboratory at the University of Pennsylvania Medical Center. CSF Aβ1-42 was measured using the Elecsys β-amyloid(1-42) assay.

### Extracellular vesicle isolation

The isolation of EVs from the plasma as well as purity validation by stochastic optical reconstruction microscopy, electron microscopy and nanoparticle tracking analysis has been described in detail elsewhere ^23^. A rabbit anti-Aβ1-42 monoclonal antibody (700254, Thermo Fisher Scientific) was labeled with the Zenon Alexa Fluor 647 Rabbit IgG labeling kit and the samples were analyzed using an Apogee Micro-PLUS flow cytometer (Apogee Flow Systems) as previously described in detail ^23^.

### 7K SomaScan Analysis

The detailed methodology can be found in the ADNI documentation (adni.loni.usc.edu) and the original publication ^39^. We used the relative fluorescence units after quality control. No longitudinal SomaScan data was available.

### Neuroimaging acquisition and processing

Detailed descriptions of ADNI neuroimaging acquisition and pre-processing can be found elsewhere (http://adni.loni.usc.edu/datasamples/pet/). We used the neocortical [^18^F]Florbetapir SUVR as well as the Desikan-Killiany-Tourville (DKT) cortical parcellation generated with FreeSurfer ^52^ that was normalized to the whole cerebellum. We only included data that passed the quality control.

### Biomarker status and staging

Participants were classified according to the amyloid/tau/neurodegeneration (A/T/N) framework using published neocortical [^18^F]Florbetapir (1.08) ^53^ or, if no amyloid-PET was available, CSF Aβ1-42 (981 pg/mL) ^54^ cut-offs for amyloid-positivity and a published CSF p-tau181 (23 pg/mL) ^55^ cut-off for tau-positivity. For N-status, we used hippocampal volume loss > 2.5 standard deviations of the mean of the 20 youngest, CU A- ADNI participants.

### Statistical analysis

All analyses were performed in R (version 4.4.1). The following packages were used for analyses and visualizations: tidyverse ^56^, tidyplots ^57^, lme4 ^58^, clusterProfiler ^59^, effectsize, broom, ggseg. To account for the amount/severity of intracerebral Aβ pathology, we calculated the ratio between the proportion of Aβ1-42 positive EVs and Aβ1-42 in the CSF (rAβ42). Pairwise comparisons between the diagnostic groups or A/T/N stages were performed using unpaired non-parametric Mann-Whitney-U tests with FDR-correction for multiple comparisons. Associations were quantified using Spearman correlation. We calculated linear models of rAβ42 with [^18^F]Florbetapir SUVR in the different cortical DKT regions and used age, sex at birth, and APOE4 status as covariates. We generated FDR-corrected *P*-values and standardized t-values for the respective DKT regions, the standardized t-values were visualized when statistical significance was reached. This was performed for all participants or separately in A+ and A-. Furthermore, the participants were separated into the highest 25% rAβ42 (rAβ42 high) and the rest (rAβ42 low). Using this dichotomization, we performed longitudinal analyses using mixed linear effects models. We tested the interaction between the two rAβ42 groups and the longitudinal trajectories of the respective other biomarkers. Age, sex at birth, APOE4 carriership, CSF Aβ1-42 and CSF p-tau181 were used as covariates. For the SomaScan data, we generated z-scores for each protein using CU A- as reference. Since we were interested in AD related pathologies, we selected the A+ participants and performed a linear model with the respective z-scores and rAβ42 and included age and sex at birth as covariates. We performed a gene ontology (GO)^60^ analysis of all proteins that were positively associated with rAβ42. We then used the proteins that defined the ”type II interferon response” GO term, were available in the 7K SomaScan data, and were significantly associated with the rAβ42 and generated a composite score as previously described ^27^ by averaging the z-scores of the respective proteins per individual. The following proteins were used: PGLYRP3, EBI3, TLR3, CEBPG, SLAMF6, LILRB4, CD2, IL1RL1, IL12B/IL23A, IL-18 Ra, TMEM106A. We then performed Spearman correlation analysis with the rAβ42 in the A+ participants. A *P*-value < 0.05 was considered statistically significant.

## Author contributions

MSW designed and conceptualized the study. LR and MSW performed most analyses and visualizations. KG, BP, CM, AH, ES, MK assisted with formal analysis and data interpretation. JZ, MS provided and generated the EV data. LR, TM, JG, TAP. SG, MG, MS, JZ, and PRN helped with data interpretation. MSW supervised the study. LR and MSW wrote the initial version of the study. All authors contributed and agreed to the final version of the study.

## Acknowledgements

Data used in preparation of this article were obtained from the Alzheimer’s Disease Neuroimaging Initiative (ADNI) database (adni.loni.usc.edu). As such, the investigators within the ADNI contributed to the design and implementation of ADNI and/or provided data but did not participate in analysis or writing of this report. A complete listing of ADNI investigators can be found at: http://adni.loni.usc.edu/wp-content/uploads/how_to_apply/ADNI_Acknowledgement_List.pdf. We thank the ADNI participants and their families who made this study possible.

## Data availability

All data used for the study can be accessed at the Alzheimer’s Disease Neuroimaging Initiative (ADNI) database (adni.loni.usc.edu). No original data was generated in the study. All code to reproduce the results are available from the corresponding author upon reasonable request.

## Conflict of interest

MSW received honoraria from Lilly for educational lectures outside the scope of this manuscript. MK received honoraria from Lilly outside the scope of this work. SG serves on scientific advisory boards for Alzeon, AmyriAD, Advantage, Eisai Canada, Enigma USA, Lilly Canada, Medesis, Lundbeck Foundation, Novo-Nordisk Canada, Okutsa, and TauRx outside the scope of this manuscript. GT has received honoraria as consultant or lecturer from Acandis, Bayer, Boehringer Ingelheim, BMS/Pfizer, Lilly, and TarGED.

## Funding

MSW is funded by the Else-Kröner-Fresenius Foundation (2023_EKMS.03), Corona Foundation (S0199/10110/2025), and German Research Foundation (DFG, WO 2835/1-1). PRN is supported by the Weston Brain Institute, Canadian Institutes of Health Research (CIHR) [MOP-11-51-31; RFN 152985, 159815, 162303], Canadian Consortium of Neurodegeneration and Aging (CCNA; MOP-1151-31 -team 1), the Alzheimer’s Association [NIRG-12-92090, NIRP-12-259245], Brain Canada Foundation (CFI Project 34874; 33397), the Fonds de Recherche du Québec – Santé (FRQS; Chercheur Boursier, 2020-VICO-279314) and the Colin J. Adair Charitable Foundation. MG is supported by the German Research Foundation (DFG, FOR 5705, Project number 523862973).

---

**Extended Data Fig. 1.**
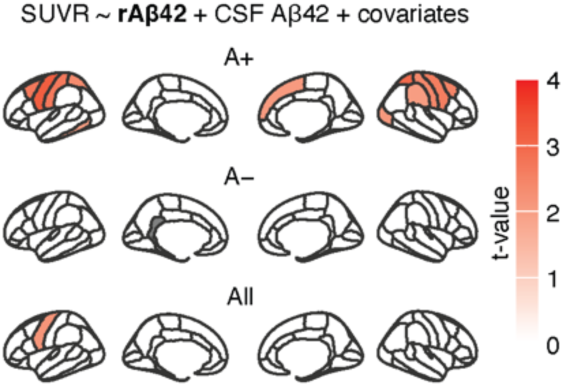
The EV/CSF Aβ42-ratio is associated with neocortical Aβ accumulation after correction for CSF Aβ42. Region-wise linear models between [^18^F]Florbetapir and rAβ42. The Desikan-Killiany-Tourville cortical parcellation was used. Age, sex at birth, CSF Aβ42 and APOE4 carriership were used as covariates. T-values are only shown where the FDR-corrected *P*-values reached statistical significance *P* < 0.05. rAβ42 = ratio between the % of Aβ42-positive plasma EVs and CSF Aβ42.

**Extended Data Fig. 2.**
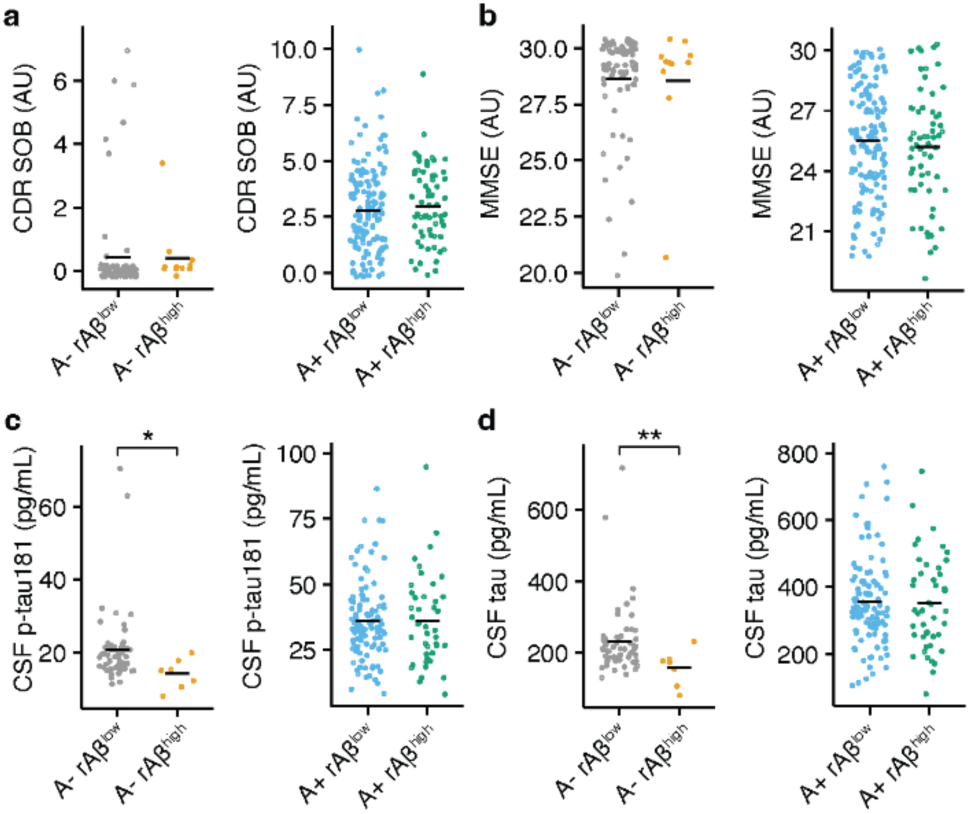
Aβ42-high with amyloid pathology have similar cognitive status and tau levels as Aβ42-low participants. **(a-d)** CDR-SOB (a), MMSE (c), CSF p-tau181 (c), and CSF total tau (d) in rAβ42 high vs. rAβ42 low people with (A+) or without (A-) Aβ pathology (A- rAβ42 low, n = 56; A+ rAβ42 low, n = 104; A- rAβ42 high, n = 7; A+ rAβ42 high, n = 45). rAβ42 = ratio between the % of Aβ42-positive plasma EVs and CSF Aβ42. Mann-Whitney-U tests were used. \**P* < 0.05, \*\**P* < 0.01.

**Extended Data Fig. 3.**
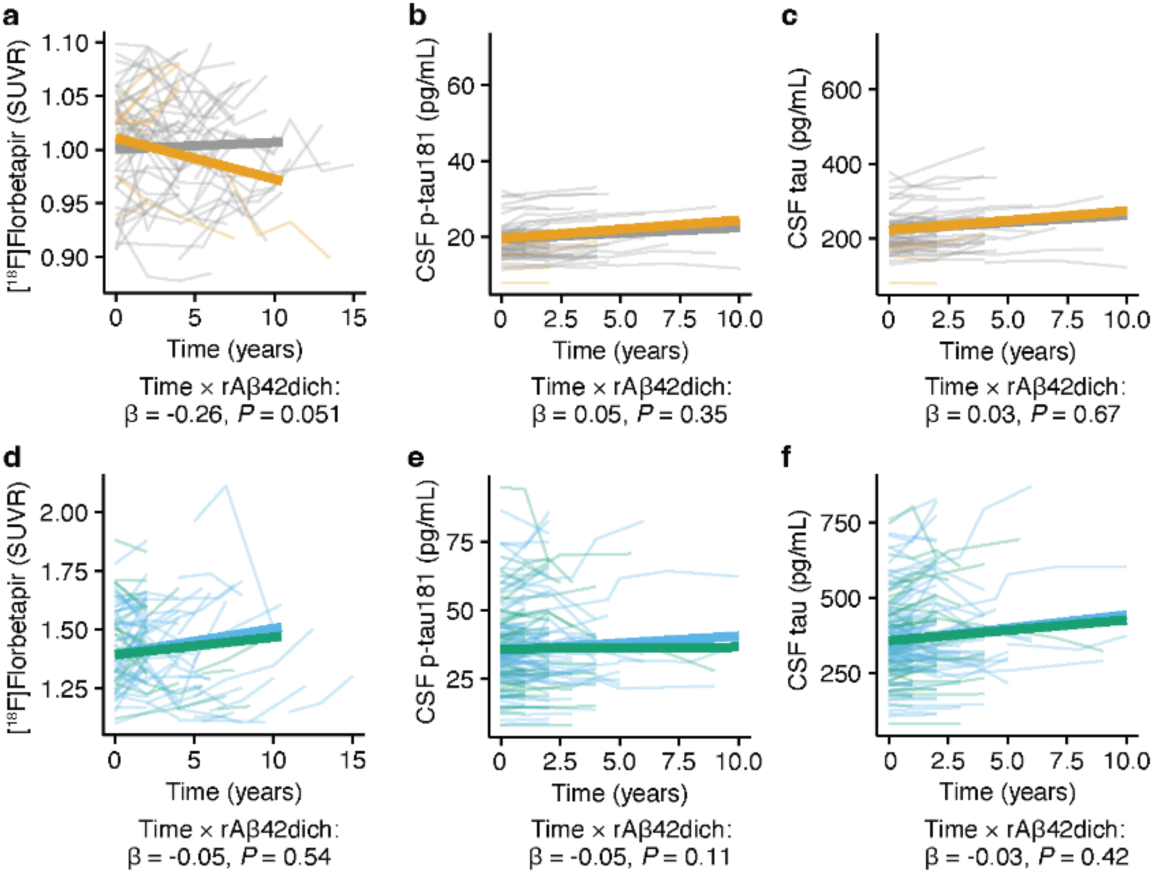
The EV/CSF Aβ42-ratio does not predict amyloid accumulation or tau increase in participants with amyloid pathology. **(a-c)** Longitudinal analysis of [^18^F]Florbetapir (a), CSF p-tau181 (b) and CSF total tau (c) in rAβ42 high and rAβ42 low participants without Aβ pathology. The standardized β and *P*-value for the interaction between time and rAβ42-status are shown in the figure. **(d-f)** Longitudinal analysis of [^18^F]Florbetapir (d), CSF p-tau181 (e) and CSF total tau (f) in rAβ42 high and rAβ42 low participants with Aβ pathology. The standardized β and *P*-value for the interaction between time and rAβ42-status are shown in the figure. Age, sex at birth, APOE4 carriership, CSF Aβ42 baseline and CSF p-tau181 baseline values were used as covariates. The respective numbers of individuals for the different groups are shown in the **Extended Data Table 1**. The full statistical results for the interaction analysis between the time and rAβ42-status are shown in **Table 2 and 3**. Statistical results of the longitudinal biomarker progression in the rAβ42 high and rAβ42 low are shown in the **Extended Data Table 2**. rAβ42 = ratio between the % of Aβ42-positive plasma EVs and CSF Aβ42.

**Extended Data Fig. 4.**
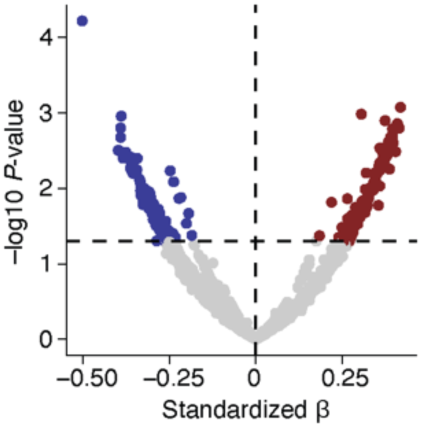
No biological pathways were enriched in the CSF of A- rAβ42-high in comparison to Aβ42-low participants. CSF 7K SomaScan data of 61 participants without amyloid pathology and available plasma EV and CSF Aβ42 was analyzed. A linear model testing the association between rAβ42 and each protein was used with age and sex at birth as covariates. The standardized β and the *P*-values are shown. Red and blue colors indicate significant up- or downregulation.

**Extended Data Table 1.** Number of participants with longitudinal data.

| Parameter | Amyloid status | rAβ42 groups | N | Follow up, years, mean (SD) | Age, years, mean (SD) |
| --- | --- | --- | --- | --- | --- |
| [<sup>18</sup>F]Florbetapir | A- | Low | 57 | 6.2 (3.2) | 73.9 (6.2) |
| [<sup>18</sup>F]Florbetapir | A- | High | 10 | 4.8 (3.7) | 73.6 (5.3) |
| CDR-SOB | A- | Low | 84 | 5.3 (3.3) | 74.2 (6.2) |
| CDR-SOB | A- | High | 13 | 5 (3.6) | 73.7 (5.4) |
| Hippocampus volume | A- | Low | 80 | 5.1 (2.8) | 74.1 (6.4) |
| Hippocampus volume | A- | High | 12 | 4.9 (3.9) | 73.2 (5.5) |
| MMSE | A- | Low | 84 | 5.4 (3.3) | 74.2 (6.2) |
| MMSE | A- | High | 13 | 5 (3.6) | 73.7 (5.4) |
| CSF p-tau181 | A- | Low | 60 | 3.3 (2.1) | 74.8 (6.1) |
| CSF p-tau181 | A- | High | 8 | 2.5 (1.2) | 73.7 (6.6) |
| CSF tau | A- | Low | 60 | 3.3 (2.1) | 74.8 (6.1) |
| CSF tau | A- | High | 8 | 2.5 (1.2) | 73.7 (6.6) |
| [<sup>18</sup>F]Florbetapir | A+ | Low | 66 | 4.9 (3.1) | 71.5 (6.9) |
| [<sup>18</sup>F]Florbetapir | A+ | High | 21 | 3.6 (2.5) | 74.3 (6.1) |
| CDR-SOB | A+ | Low | 156 | 3.4 (2.8) | 73.5 (7.1) |
| CDR-SOB | A+ | High | 64 | 2.5 (2) | 74.7 (7.4) |
| Hippocampus volume | A+ | Low | 135 | 3.2 (2.3) | 72.8 (7.2) |
| Hippocampus volume | A+ | High | 53 | 2.5 (2) | 73.7 (6.9) |
| MMSE | A+ | Low | 156 | 3.4 (2.8) | 73.4 (7.1) |
| MMSE | A+ | High | 64 | 2.5 (2) | 74.7 (7.4) |
| CSF p-tau181 | A+ | Low | 112 | 2.5 (1.8) | 73.9 (7.2) |
| CSF p-tau181 | A+ | High | 46 | 2.2 (1.6) | 74.6 (7.5) |
| CSF tau | A+ | Low | 112 | 2.5 (1.8) | 73.9 (7.2) |
| CSF tau | A+ | High | 46 | 2.2 (1.6) | 74.6 (7.5) |

**Extended Data Table 2.** Longitudinal biomarker progression in rAβ42 high/low and A+/A- individuals.

| Parameter | Amyloid status | rAβ42 groups | *P*-value | Standardized $\beta$ (95% CIs) |
| --- | --- | --- | --- | --- |
| [<sup>18</sup>F]Florbetapir | A- | Low | 0.34 | 0.04 (-0.04, 0.12) |
| [<sup>18</sup>F]Florbetapir | A- | High | 0.12 | -0.18 (-0.4, 0.05) |
| CDR-SOB | A- | Low | 0.19 | 0.03 (-0.01, 0.07) |
| CDR-SOB | A- | High | 0.96 | < 0.001 (-0.09, 0.08) |
| Hippocampus volume | A- | Low | < 0.01 | -0.24 (-0.29, -0.2) |
| Hippocampus volume | A- | High | < 0.01 | -0.21 (-0.33, -0.1) |
| MMSE | A- | Low | 0.1 | -0.06 (-0.14, 0.01) |
| MMSE | A- | High | 0.01 | 0.23 (0.06, 0.4) |
| CSF p-tau181 | A- | Low | < 0.01 | 0.09 (0.06, 0.11) |
| CSF p-tau181 | A- | High | 0.01 | 0.07 (0.02, 0.13) |
| CSF tau | A- | Low | < 0.01 | 0.11 (0.07, 0.14) |
| CSF tau | A- | High | 0.05 | 0.07 (0, 0.14) |
| [<sup>18</sup>F]Florbetapir | A+ | Low | < 0.01 | 0.15 (0.07, 0.23) |
| [<sup>18</sup>F]Florbetapir | A+ | High | 0.13 | 0.14 (-0.04, 0.32) |
| CDR-SOB | A+ | Low | < 0.01 | 0.49 (0.42, 0.57) |
| CDR-SOB | A+ | High | < 0.01 | 0.52 (0.4, 0.64) |
| Hippocampus volume | A+ | Low | < 0.01 | -0.28 (-0.32, -0.25) |
| Hippocampus volume | A+ | High | < 0.01 | -0.29 (-0.33, -0.25) |
| MMSE | A+ | Low | < 0.01 | -0.42 (-0.5, -0.34) |
| MMSE | A+ | High | < 0.01 | -0.48 (-0.6, -0.37) |
| CSF p-tau181 | A+ | Low | < 0.01 | 0.06 (0.02, 0.09) |
| CSF p-tau181 | A+ | High | 0.54 | 0.01 (-0.03, 0.06) |
| CSF tau | A+ | Low | < 0.01 | 0.12 (0.08, 0.15) |
| CSF tau | A+ | High | < 0.01 | 0.07 (0.04, 0.11) |

## References

1. Livingston, G. et al. Dementia prevention, intervention, and care: 2024 report of the Lancet standing Commission. Lancet 404, 572–628 (2024).

2. Knopman, D. S. et al. Alzheimer disease. Nat. Rev. Dis. Prim. 7, 33 (2021).

3. Heneka, M. T. et al. Neuroinflammation in Alzheimer disease. Nat. Rev. Immunol. 25, 321–352 (2025).

4. Haney, M. S. et al. APOE4/4 is linked to damaging lipid droplets in Alzheimer’s disease microglia. Nature 628, 154–161 (2024).

5. Chen, X. et al. Microglia-mediated T cell infiltration drives neurodegeneration in tauopathy. Nature 615, 668–677 (2023).

6. Bellaver, B. et al. Astrocyte reactivity influences amyloid-β effects on tau pathology in preclinical Alzheimer’s disease. Nat. Med. 29, 1775–1781 (2023).

7. Zhou, Y. et al. Human and mouse single-nucleus transcriptomics reveal TREM2-dependent and TREM2-independent cellular responses in Alzheimer’s disease. Nat. Med. 26, 131–142 (2020).

8. McAlpine, C. S. et al. Astrocytic interleukin-3 programs microglia and limits Alzheimer’s disease. Nature 595, 701–706 (2021).

9. Therriault, J. et al. Association of Phosphorylated Tau Biomarkers With Amyloid Positron Emission Tomography vs Tau Positron Emission Tomography. JAMA Neurol. 80, 188 (2023).

10. Janelidze, S. et al. Plasma Phosphorylated Tau 217 and Aβ42/40 to Predict Early Brain Aβ Accumulation in People Without Cognitive Impairment. JAMA Neurol. 81, 947 (2024).

11. Horie, K. et al. CSF MTBR-tau243 is a specific biomarker of tau tangle pathology in Alzheimer’s disease. Nat. Med. 29, 1954–1963 (2023).

12. Ashton, N. J. et al. Differential roles of Aβ42/40, p-tau231 and p-tau217 for Alzheimer’s trial selection and disease monitoring. Nat. Med. 28, 2555–2562 (2022).

13. Woo, M. S. et al. Identification of late-stage tau accumulation using plasma phospho-tau217. eBioMedicine 109, 105413 (2024).

14. Woo, M. S. et al. Plasma pTau-217 and N-terminal tau (NTA) enhance sensitivity to identify tau PET positivity in amyloid-β positive individuals. Alzheimer’s Dement. 20, 1166–1174 (2024).

15. Therriault, J. et al. Biomarker-based staging of Alzheimer disease: rationale and clinical applications. Nat. Rev. Neurol. 20, 232–244 (2024).

16. Oh, H. S.-H. et al. A cerebrospinal fluid synaptic protein biomarker for prediction of cognitive resilience versus decline in Alzheimer’s disease. Nat. Med. 31, 1592–1603 (2025).

17. Wang, Y.-T. et al. The relation of synaptic biomarkers with Aβ, tau, glial activation, and neurodegeneration in Alzheimer’s disease. Transl. Neurodegener. 13, 27 (2024).

18. van Niel, G., D’Angelo, G. & Raposo, G. Shedding light on the cell biology of extracellular vesicles. Nat. Rev. Mol. Cell Biol. 19, 213–228 (2018).

19. Chatterjee, M. et al. Plasma extracellular vesicle tau and TDP-43 as diagnostic biomarkers in FTD and ALS. Nat. Med. 30, 1771–1783 (2024).

20. Fowler, S. L. et al. Tau filaments are tethered within brain extracellular vesicles in Alzheimer’s disease. Nat. Neurosci. 28, 40–48 (2025).

21. Ruan, Z. et al. Alzheimer’s disease brain-derived extracellular vesicles spread tau pathology in interneurons. Brain 144, 288–309 (2021).

22. González-Molina, L. A. et al. Extracellular Vesicles From 3xTg-AD Mouse and Alzheimer’s Disease Patient Astrocytes Impair Neuroglial and Vascular Components. Front. Aging Neurosci. 13, (2021).

23. Tian, C. et al. Blood extracellular vesicles carrying synaptic function- and brain-related proteins as potential biomarkers for Alzheimer’s disease. Alzheimer’s Dement. 19, 909–923 (2023).

24. Vilcaes, A. A., Chanaday, N. L. & Kavalali, E. T. Interneuronal exchange and functional integration of synaptobrevin via extracellular vesicles. Neuron 109, 971–983.e5 (2021).

25. Ikezu, T., Yang, Y., Verderio, C. & Krämer-Albers, E.-M. Extracellular Vesicle-Mediated Neuron–Glia Communications in the Central Nervous System. J. Neurosci. 44, e1170242024 (2024).

26. Buzas, E. I. The roles of extracellular vesicles in the immune system. Nat. Rev. Immunol. 23, 236–250 (2023).

27. Woo, M. S. et al. Glia inflammation and cell death pathways drive disease progression in preclinical and early AD. EMBO Mol. Med. (2025) doi:10.1038/s44321-025-00316-1.

28. Esquivel, R. N. et al. Clinical validation of the Lumipulse G β-amyloid ratio (1-42/1-40) in a subset of ADNI CSF samples. Alzheimer’s Dement. 17, (2021).

29. Barthélemy, N. R. et al. Highly accurate blood test for Alzheimer’s disease is similar or superior to clinical cerebrospinal fluid tests. Nat. Med. 30, 1085–1095 (2024).

30. Han, J., Zhang, X., Kang, L. & Guan, J. Extracellular vesicles as therapeutic modulators of neuroinflammation in Alzheimer’s disease: a focus on signaling mechanisms. J. Neuroinflammation 22, 120 (2025).

31. Beretta, C. et al. Extracellular vesicles from amyloid-β exposed cell cultures induce severe dysfunction in cortical neurons. Sci. Rep. 10, 19656 (2020).

32. Cone, A. S. et al. Alix and Syntenin-1 direct amyloid precursor protein trafficking into extracellular vesicles. BMC Mol. Cell Biol. 21, 58 (2020).

33. Su, H. et al. Characterization of brain-derived extracellular vesicle lipids in Alzheimer’s disease. J. Extracell. Vesicles 10, (2021).

34. Yang, L. et al. Decoding adipose–brain crosstalk: Distinct lipid cargo in human adipose-derived extracellular vesicles modulates amyloid aggregation in Alzheimer’s disease. Alzheimer’s Dement. 21, (2025).

35. Mustapic, M., Tran, J., Craft, S. & Kapogiannis, D. Extracellular Vesicle Biomarkers Track Cognitive Changes Following Intranasal Insulin in Alzheimer’s Disease. J. Alzheimer’s Dis. 69, 489–498 (2019).

36. Sattlecker, M. et al. Alzheimer’s disease biomarker discovery using SOMAscan multiplexed protein technology. Alzheimer’s Dement. 10, 724–734 (2014).

37. Shen, Y. et al. CSF proteomics identifies early changes in autosomal dominant Alzheimer’s disease. Cell 187, 6309–6326.e15 (2024).

38. Pichet Binette, A. et al. Proteomic changes in Alzheimer’s disease associated with progressive Aβ plaque and tau tangle pathologies. Nat. Neurosci. 27, 1880–1891 (2024).

39. Ali, M. et al. Multi-cohort cerebrospinal fluid proteomics identifies robust molecular signatures across the Alzheimer disease continuum. Neuron 113, 1363–1379.e9 (2025).

40. Pascoal, T. A. et al. Microglial activation and tau propagate jointly across Braak stages. Nat. Med. 27, 1592–1599 (2021).

41. Mancuso, R. et al. Xenografted human microglia display diverse transcriptomic states in response to Alzheimer’s disease-related amyloid-β pathology. Nat. Neurosci. 27, 886–900 (2024).

42. Venegas, C. et al. Microglia-derived ASC specks cross-seed amyloid-β in Alzheimer’s disease. Nature 552, 355–361 (2017).

43. Jorfi, M. et al. Infiltrating CD8+ T cells exacerbate Alzheimer’s disease pathology in a 3D human neuroimmune axis model. Nat. Neurosci. 26, 1489–1504 (2023).

44. Woo, M. S. et al. STING orchestrates the neuronal inflammatory stress response in multiple sclerosis. Cell 187, 4043–4060.e30 (2024).

45. Woo, M. S. et al. The immunoproteasome disturbs neuronal metabolism and drives neurodegeneration in multiple sclerosis. Cell 188, 4567–4585.e32 (2025).

46. Arvanitaki, E. S. et al. Microglia-derived extracellular vesicles trigger age-related neurodegeneration upon DNA damage. Proc. Natl. Acad. Sci. 121, (2024).

47. Zhang, L. et al. Extracellular vesicles from hypoxia-preconditioned microglia promote angiogenesis and repress apoptosis in stroke mice via the TGF-β/Smad2/3 pathway. Cell Death Dis. 12, 1068 (2021).

48. De Paula, G. C. et al. Extracellular vesicles released from microglia after palmitate exposure impact brain function. J. Neuroinflammation 21, 173 (2024).

49. Brenna, S. et al. Characterization of brain-derived extracellular vesicles reveals changes in cellular origin after stroke and enrichment of the prion protein with a potential role in cellular uptake. J. Extracell. Vesicles 9, (2020).

50. Gui, Y. et al. Cystatin C loaded in brain-derived extracellular vesicles rescues synapses after ischemic insult in vitro and in vivo. Cell. Mol. Life Sci. 81, 224 (2024).

51. Cleary, J. A., Kumar, A., Craft, S. & Deep, G. Neuron-derived extracellular vesicles as a liquid biopsy for brain insulin dysregulation in Alzheimer’s disease and related disorders. Alzheimer’s Dement. 21, (2025).

52. Fischl, B. FreeSurfer. Neuroimage 62, 774–781 (2012).

53. Landau, S. M. et al. Amyloid-β Imaging with Pittsburgh Compound B and Florbetapir: Comparing Radiotracers and Quantification Methods. J. Nucl. Med. 54, 70–77 (2013).

54. Dumurgier, J. et al. A Pragmatic, Data-Driven Method to Determine Cutoffs for CSF Biomarkers of Alzheimer Disease Based on Validation Against PET Imaging. Neurology 99, (2022).

55. Tan, M.-S. et al. Longitudinal trajectories of Alzheimer’s ATN biomarkers in elderly persons without dementia. Alzheimers. Res. Ther. 12, 55 (2020).

56. Wickham, H. et al. Welcome to the Tidyverse. J. Open Source Softw. 4, 1686 (2019).

57. Engler, J. B. Tidyplots empowers life scientists with easy code-based data visualization. iMeta (2025) doi:10.1002/imt2.70018.

58. Bates, D., Mächler, M., Bolker, B. & Walker, S. Fitting Linear Mixed-Effects Models Using lme4. J. Stat. Softw. 67, (2015).

59. Yu, G., Wang, L.-G., Han, Y. & He, Q.-Y. clusterProfiler: an R Package for Comparing Biological Themes Among Gene Clusters. Omi. A J. Integr. Biol. 16, 284–287 (2012).

60. Ashburner, M. et al. Gene Ontology: tool for the unification of biology. Nat. Genet. 25, 25–29 (2000).

